# Family lexicon: using language models to encode memories of personally familiar and famous people and places in the brain

**DOI:** 10.1101/2023.08.23.554436

**Authors:** Andrea Bruera, Massimo Poesio

## Abstract

Knowledge about personally familiar people and places is extremely rich and varied, involving pieces of semantic information connected in unpredictable ways through past autobiographical memories. In this work we investigate whether we can capture brain processing of personally familiar people and places using subject-specific memories, after transforming them into vectorial semantic representations using language models. First we asked participants to provide us with the names of the closest people and places in their lives. Then we collected open-ended answers to a questionnaire, aimed at capturing various facets of declarative knowledge. We collected EEG data from the same participants while they were reading the names and subsequently mentally visualizing their referents. As a control set of stimuli, we also recorded evoked responses to a matched set of famous people and places. We then created original semantic representations for the individual entities using language models. For personally familiar entities, we used the text of the answers to the questionnaire. For famous entities, we employed their Wikipedia page, which reflects shared declarative knowledge about them. Through whole-scalp time-resolved and searchlight encoding analyses we found that we could capture how the brain processes one’s closest people and places using person-specific answers to questionnaires, as well as famous entities. Encoding performance was significant in a large time window (200-800ms). In terms of spatio-temporal clusters, two main axes where encoding scores are significant emerged, in bilateral temporo-parietal electrodes first (200-500ms) and frontal and posterior central electrodes later (500-700ms). We also found that XLM, a contextualized language model or large language model, provided superior encoding scores when compared with a simpler static language model as word2vec. Overall, these results indicate that language models can capture subject-specific semantic representations as they are processed in the human brain, by exploiting small-scale distributional lexical data.

> *My parents had five children. We now live in different cities, some of us in foreign countries, and we don’t write to each other often. When we do meet up we can be indifferent or distracted. But for us it takes just one word. It takes one word, one sentence, one of the old ones from our childhood, heard and repeated countless times. All it takes is for one of us to say “We haven’t come to Bergamo on a military campaign,” or “Sulfuric acid stinks of fart,” and we immediately fall back into our old relationships, our childhood, our youth, all inextricably linked to those words and phrases*.
>
> excerpt from Family Lexicon, by Natalia Ginzburg

## Introduction

Being asked to describe one’s closest friend, or one’s favourite neighbourhood, is not an easy question to answer. One will find that just describing physical and personality traits is not enough - to do justice to that specific person or place it will be necessary to bring up much more. Anecdotes, past events and stories involving them, together with other disparate pieces of information that just come up to one’s mind when talking about familiar entities. And, taken together, this will form an extremely idiosyncratic mixture, that however captures the fundamental uniqueness of that person or place in one’s memory.

In cognitive neuroscience, as a reflection of this, it has been found that knowledge for individual entities, such as people and places, is particularly rich and multifaceted [1–3]. Aside from its episodic (event-specific) and semantic (encyclopedic) components [4–6], it seems to strongly involve another type of knowledge, called ‘personal semantics’ [7]. Personal semantics can be described as a type of knowledge which is abstracted from individual events and occurrences, but not fully so. An example could be knowing what a friend enjoys doing. It is not entirely encyclopedic, as it is dependent from memories of repeated events that took place in the past. Nor is it completely episodic, as it does not strictly depend from individual instances of that event. In this sense, is part of personal semantics.

Episodic, semantic and personal knowledge compose declarative, or explicit, memory [6, 7]. By definition, declarative memory is knowledge that can be expressed through natural language [8].

For generic concepts, such as a cello, declarative knowledge is widely shared across a community of speakers. Because of this, it can be easily extracted in a data-driven fashion from large collections of text (called **corpora**) such as Wikipedia. This is done by creating vectorial representations reflecting distributional properties of their corresponding words in the corpora through **language models**, also called a **distributional semantics model**. Such approaches follow the hunch that the semantic content of a word can be captured by the way in which this word is used [9]. This, in turn, can be operationalized as the so-called distributional hypothesis - that words having similar meanings will be found in similar linguistic contexts [10]. The resulting representations are traditionally called **word vectors**. Word vectors have been shown to capture and encompass in a single high-dimensional vectorial space multiple traditional semantic dimensions proposed in cognitive science [11–14]. This can explain their effectiveness at modelling semantic processing, both in behaviour [15, 16] and in the brain [17–20].

However, when it comes to personally familiar people or places, two main challenges arise. First, the extraction of semantic representations for concepts based on distributional information requires words to be frequent enough in the corpora in order to robustly capture their meaning [21–25]. Despite their fundamental importance in our lives, personally familiar people and places (a close friend, one’s favourite neighbourhood) never - or extremely rarely - get mentioned in large-scale corpora such as Wikipedia. Therefore, it seems impossible, in principle, to capture their meaning by way of word vectors.

Secondly, even if one were able to sidestep this issue, a more pervasive one would emerge: namely, that each person has highly idiosyncratic and subjective ways of perceiving and describing personally familiar people and places. This makes it hard to capture semantic representations from recollections of autobiographic memories expressed in natural language, which constitute an exceptionally diverse and reduced linguistic dataset [26–28].

Such limitations posed by language models have had an impact on studies employing them as models to capture semantic representations in the brain. As a consequence of the need for sufficient training data, previous work has focused on generic concepts for whom a representation could be obtained from large corpora [17, 18] – or, in the case of individual entities, on famous entities which are mentioned with enough frequency in corpora [29]. The only partial exception taking into consideration subject-specific semantic knowledge is, to our knowledge, [30], which however focused on personal interpretations of generic concepts and not of individual entities. Previous work in cognitive neuroscience looking at subject-specific knowledge of individual entities has not used distributional linguistic information, but rather semantic dimensions defined a priori by the experimenters [31–33].

In this work we set out to investigate whether we could build unique vectorial semantic representations for personally familiar entities, such as people and places, that could encode the way in which the brain of each subject processes such entities. Our approach follows a framework that has been recently emerging in neuroscience, aiming at recognizing and effectively accounting for the uniqueness of individuals and of their cognitive and neural processes [30, 34–36].

In our experiment, first we captured person-specific knowledge in the form of text. We did so by asking subjects to talk about the most important people and places in their lives (eight people and eight places; see Fig 1). This allowed us to collect textual data regarding semantic knowledge (physical and personality traits), episodic memory (the most salient memories involving a person/place) and personal semantics knowledge (operationalized as words and topics associated with each person/place). We then used language models to encode each subject’s text, thus creating subject-specific vector representations of personally familiar entities from small-scale textual distributional information. We also created in a mirrored way vector representations for eight famous people and eight famous places, as a ‘control’ set of entities, for which it is known that language models are able to create reliable semantic representations given their high frequency in corpora [29, 37, 38].

In parallel, we collected electroencephalography (EEG) data while the same subjects were reading the entities’ names (both famous and familiar). We ran a set of encoding analyses, looking at where in time (time-resolved encoding using all electrodes) and in space (spatio-temporal searchlight encoding, looking at clusters of electrodes separately) our subject-specific vector representations captured brain processing. Names were used instead of faces in order to avoid non-semantic, low-level visual differences among different categories of stimuli (people and places, famous and personally familiar).

We compared two language models for Italian, the language in which the experiment was carried out, from two different families of language models. The first model is **word2vec** [39], a so-called static model. In static models, each word is represented as a single, fixed vector representation. This captures distributional information encoded in the training data only for word types (i.e. all aggregated mentions of words), but not tokens (i.e. actual individual occurrences in context). Static models are the most commonly used models in brain encoding/decoding [20]. The second model is XLM-Roberta-large (**XLM**, [40]), a so-called **contextualized model**, or **large language model** (**LLM**). LLMs are more recent and complex models, based on the Transformer architecture [41]. They do not represent word *types* (as static models do), but word *tokens*: given a linguistic context, such as a sentence, the vectors for the words are created by adapting pre-trained representations to the specific sentence - thus incorporating both type- and token-level information. This type of model has never been tested, to our knowledge, as a model for personal, subject-specific semantic representations. We used XLM because it is a state-of-the-art multilingual model which can therefore be used to encode text in Italian.

The results reported in Fig s 3-4 indicate that by encoding personal memories with language models it was possible to create semantic representations of individuals that captured the way in which they are processed in the brain. Importantly, we were able to create semantic representations for the most important people and places in each subject’s life using only the reduced amount of text available to us through a questionnaire –i.e. small-scale distributional lexical information. Furthermore, this approach worked also for a matched set of famous entities, for which non-personal textual information obtained from Wikipedia is used.

## Methods

### Stimuli

**Fig 1.**
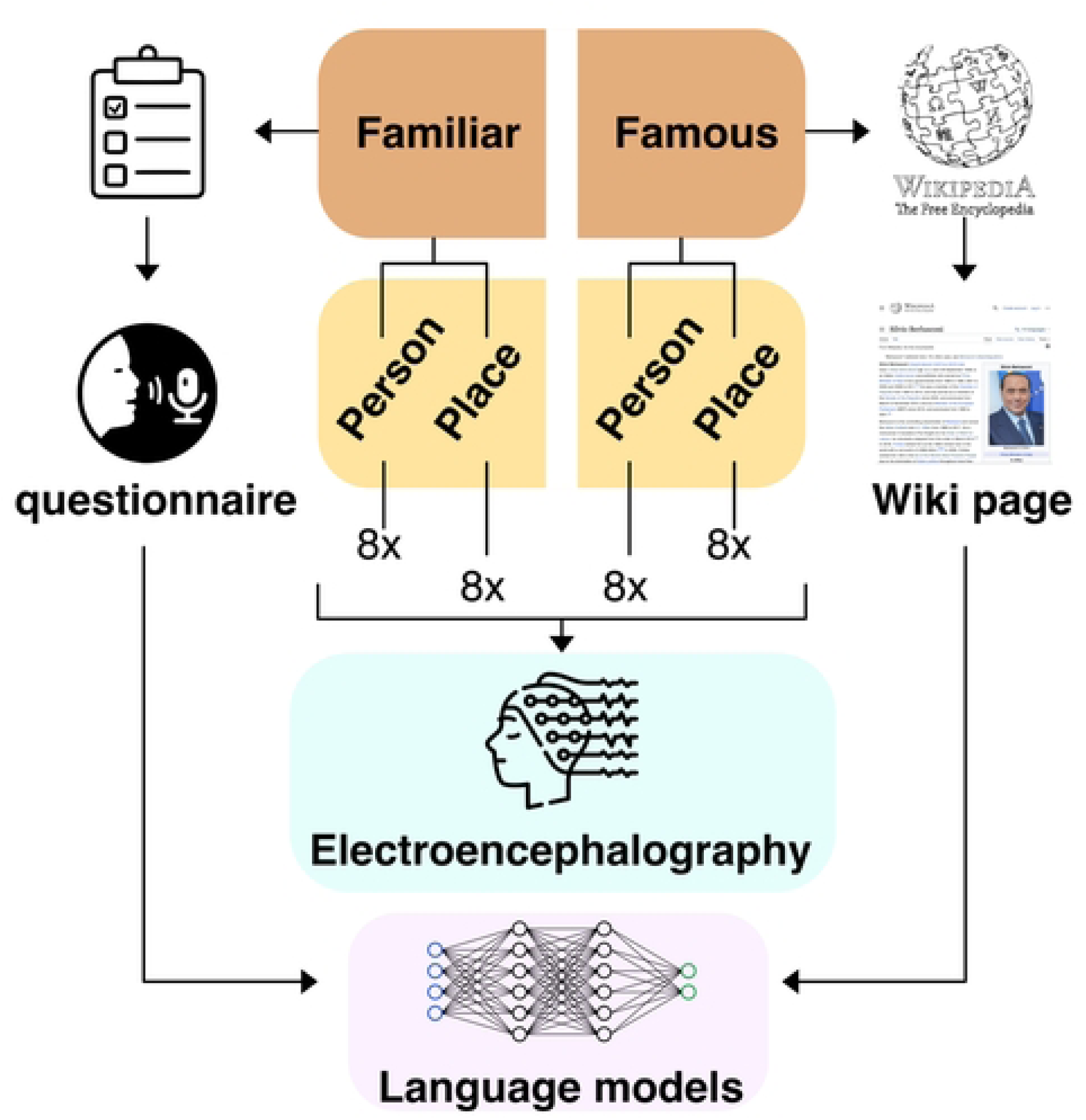
Visualization of the experimental setup. Our stimuli were divided into proper and personally familiar names. For each level of familiarity, sixteen famous individual entities (eight people and eight places) were present. Famous entities were selected after controlling for familiarity, imageability and name length. Personally familiar names were obtained by asking each participant to provide the names themselves. We also collected short amounts of text describing each entity (Wikipedia pages for famous entities, and answers to a questionnaire for personally familiar people and places). From these, we extracted one semantic vector per entity using a language model. Then we collected the EEG data, and carried out separately for each participant the encoding analyses.

### Famous entities

As experimental stimuli, we selected eight famous people and eight famous places to be used for all participants (see Fig 1, left portion). Before each EEG experimental session took place, We made sure that subjects had not either personally known any famous person or visited a famous place among the stimuli. Only two subjects had visited one of the famous places selected as experimental stimuli. In those cases we substituted the names with other place names matched for length, familiarity and imageability.

We balanced the stimuli in terms of name length, familiarity and imageability, in order to avoid significant differences across the two semantic categories. The final set of stimuli was selected from a larger set of 100 stimuli, for which we obtained familiarity and imageability ratings. The subjects for the rating tasks (carried out separately; 33 subjects for familiarity, 30 for imageability) were native speakers of Italian, and none of them took part to the EEG experiment. Familiarity was defined in the same way as in [42] - a quantification on a scale from 1 to 5 of the number of cumulative encounters with an individual entity, across time and media. The average familiarity overall was 3.59; average familiarity for people was 3.74, while that for places was 3.44, and their difference across people and places was not statistically significant (*t* = 1.63, *p* = 0.12). Imageability was defined after [43] as the ease or difficulty of arousing a mental image. Imageability was controlled because, during the experiment, subjects were asked to read names and picture their referents mentally (see Section). We used the most common scale in imageability rating experiments, going from 1 to 7 [44]. The average imageability across all entities was 4.97. Average imageability for people was 5.17, while average imageability for places was 4.78. The difference between the two was not statistically significant (*t* = 0.88, *p* = 0.39). Name length was on average overall 13.5; average person name length was 14.1, while average place name length was 13, and the difference across categories was again not significant (*t* = 0.68, *p* = 0.5). All ratings are provided together with the code for replication.

For each famous individual entity we also collected the text from their Wikipedia page, under the assumption that such texts are a source of explicit knowledge regarding individual entities that can be mapped to the brain using their distributional information [45]. These texts will be used as described below in order to extract semantic representations using language models.

### Personally familiar entities

Before starting the EEG experiment, we asked participants to provide the names for eight people and eight places (see Fig 1, left portion). As a framework, we followed research on social circles [46]. For people we focused, in our definition, on members of the so-called ‘support clique’. This circle consists of people with whom one has a positive relationship, is in touch regularly, and from whom one would seek personal advice or help [47]. For places, we tried to match as closely as possible the definition given for people, since no relevant literature was available. We defined ‘support places’ as places with whom one has a special, positive relationship, and they are places where one would (if possible) return to when in a situation of distress. We provide in the code the text of the specific instructions given to the subjects, translated to English. Notice that participants were not only asked to provide names, but also to provide either the person’s occupation or the type of place (i.e. monument, city, river, etc.), to be used during the experimental paradigm (see Section).

Additionally, subjects were asked to respond to a questionnaire, whose aim was capturing the main components of declarative knowledge about personally familiar entities – i.e. what is explicitly known about them. We looked at the two components most traditionally associated with explicit knowledge (semantic memory and episodic memory, which differ by being respectively dependent or not from specific events [4, 6]), as well as at what has been called personal semantics [7], which is a highly personal type of knowledge, lying at the intersection of episodic and semantic memory (see Introduction). The questionnaire therefore involved nine questions, divided equally among the three types of knowledge. For semantic memory, we asked to talk about the type of relationship being shared, and physical and personality traits; for episodic memory, we asked to talk about how each entity was first encountered, as well as two most recent salient autobiographical episodes involving that entity; for personal semantics, we asked to name up to 10 words that came to mind when thinking about that entity, what one would talk about (for people) or do (for places) if they met or visited that entity, and finally a sentence that is associated with a person/place. We recorded the answers using a Zoom H2 stereo digital audio recorder, and we automatically transcribed the texts using Microsoft Azure’s Speech-To-Text service^1^. Before extracting the semantic representations from the texts using the language models (see Section), we checked the automatic transcriptions in order to verify that quality was sufficiently good, which we found to be the case.

In the case of personally familiar names, we could not control in advance for name length, since participants came up with the names. Therefore, we implement in the analyses a procedure for explicitly removing all variance associated with name length from the EEG data, described below in Section.

Notice that, because of privacy reasons, it is not possible to publicly share the responses to the questionnaires, nor the subject-specific computational representations extracted from them. The parts of the data that could be published, together with the code, are publicly available online, on the Open Science Foundation website (https://osf.io/sjtmn/?view_only=49dcdbf7aa2649fa9e376f07c26ee417). The same dataset was used for a different study, currently under review [48].

### Experimental paradigm

Thirty-tree right-handed subjects (age from 20 to 31 years old, with 21 female participants) took part to the experiment. Sample size was determined following [49], where the authors show thirty-two subjects is an adequate sample size for event-related potentials (**ERP**) studies. As the experiment was conducted in Italian, all the subjects were native Italian speakers. All experimental procedures were approved by the Ethical Committee of SISSA, Trieste, where the data were collected, and subjects gave their written informed consent.

Before the EEG experiment subjects provided names for personally familiar stimuli, as well their occupations or types of places. Then, participants took part to 24 experimental EEG runs. Each name would appear once per run, in randomized order. For each name we thus recorded twenty-four ERPs, which were averaged after preprocessing and before entering the analyses. This is routinely done in encoding/decoding studies for EEG in order to improve the signal-to-noise ratio [50].

Each trial proceeded as follow. First, a fixation cross appeared for 500 ms. Then a name appeared on screen for 500 milliseconds, followed by a fixation cross lasting 1 second. Subject were instructed to read the name and mentally picture its referent. This was done in order to elicit semantic processing. Also, mental imagery task has been shown to provide good performance in encoding/decoding [18, 29, 51]. After the fixation cross disappeared, a binary yes/no question appeared on screen. The question was added exclusively to ensure participants actually engaged in the task. To avoid strategic preparation, questions were randomized among two templates. They could reflect either a coarse-category question (e.g. ‘is it a person/place?’), or a fine-grained question (e.g. ‘does the name refer to a student?’ or ‘does the name refer to a city?’), a methodology previously used in [29, 52]. Questions were balanced between yes/no answers, and subjects could answer using two keys. In the case of fine-grained questions, also the occupation or place type was randomized.

### EEG data collection and preprocessing

The EEG data was collected using a BIOSEMI ActiveTwo system with 128 channels, recording unfiltered signals at a sampling rate of 2048 hz. We also collected signals from two electro-oculogram channels (EOG) so as to be able to use them later for artifact rejection with Indipendent Component Analysis (ICA; details below). For preprocessing, we adapted an automated procedure using MNE [53], previously validated in [54].

We set the montage to the default for the 128-channel BIOSEMI system. A visualization of the montage, together with the codes for the channels, is included in the publicly available code and can be retrieved at https://www.biosemi.com/pics/cap_128_layout_medium.jpg. Then, following [54], we used the standard ICA-based ocular artifact removal implemented in MNE. We applied a low-pass filter to the ERP data to 80 hz [55]. Then we epoched the data and subsampled it to 256hz, and removed the independent components correlating the most with eye-movement artifacts [54]. Baseline correction consisted of the average signal between 100 and 0ms before the appearance of a stimulus. We then used the autoreject algorithm [56] to interpolate bad channels, and we set the reference to the grand average.

The final preprocessing step was removing all the variance associated with word length from the EEG signal using cross-validated confound regression, which was validated in [57, 58] and whose implementation is publicly available.

### Models

#### Language models

In order to create semantic representations for famous and personally familiar entities, we employed the texts described in Section as inputs for two types of language models, one static (word2vec) and one contextualized (XLM-Roberta-large). For famous entities we used the Wikipedia pages, for personally familiar entities the transcriptions of the answers to the questionnaire. The rationale for this procedure is that words appearing in an entity’s text (its distributional lexical information) carry semantic information with respect to that entity.

The procedure was matched across models, so as to avoid methodological confounds. For each individual entity, we retained all content (open-class) words from the corresponding descriptive text (Wikipedia or questionnaire), and we obtained an entity representation by averaging the vectors for those words. Notice that we excluded closed-class words such as function words since they do not carry semantic information, and in static models they are hard to represent properly [39, 59, 60]. We combine words vectors by averaging since this procedure has been previously validated as a way to combine word vectors that captures meaning both in the brain [30, 45] and in downstream NLP tasks [61].

At the end of the procedure, we obtained one vector representation for each individual entity per model, capturing the distributional lexical information contained in our small-scale textual data.

Word2vec is a feedforward neural network which learns vector representations (word vectors) from large-scale corpora [39]. Therefore, one single vector representation is created for each word type, regardless of homonymy and polysemy phenomena (e.g. the word ‘bat’ is modelled as a single vector, collapsing the two senses of animal and baseball instrument in a single representation). Such vector representations have been interpreted as models of cognitive semantic representations [11, 23, 62] and have been shown to capture well lexical processing [15, 16, 45].

We pre-trained a word2vec model on the Italian version of Wikipedia using parameters suggested in the literature. We used a skip-gram training regime, a window size of 10 words and a vector size of 300 dimensions [15, 63].

A contextualized language model [61], also often called a large language model (LLM), is a deep neural network based on the Transformer architecture [41]. Its main difference with respect to a static language model is that, by design, it is aimed at the representation of words in context – i.e. sentences, paragraphs or longer passages. In LLMs, there are no ‘static’ word representations, but instead representations are adapted to each linguistic context used as input. This allows to capture fine-grained shifts in meaning, both at at a lexical and at supra-lexical level (e.g. discourse, dialogue). Notice, however, that this is achieved at the cost of an extremely more sophisticated neural network architecture and costly training procedure [64]. Since publicly available LLMs for languages other than English are not on a par in terms of quality with their English counterpart [65], and our questionnaires were collected in Italian, we used a state-of-the-art multi-language model, XLM-Roberta-large [40]. Multi-language models are not specialized for individual languages. They are particularly effective as they exploit transfer of cross-lingual semantic information during pre-training [66, 67].

Since LLMs are designed to take as input natural language sentences, we used XLM to encode full sentences - but we averaged only the vectors corresponding to content words (see Section). Following previous work, we modelled entity representations using so-called (sentence) representation pooling for the top four layer [37, 61] – which consists of averaging the chosen words in each sentence, and then averaging the vectors obtained for the individual sentences. In order to ensure that text portions were long enough to work well with XLM, which has been shown to work better with rather long passages [68], if a sentence were shorter than 20 words (the most common sentence length in English is between 10 and 30 words [69]) we would unite it with the following sentence; and so on until a passage of at least 20 words was created.

Each entity representation in XLM was represented by a single vector having 1024 dimensions.

#### Non-semantic baseline models

We also report time-resolved encoding scores for two baseline, non-semantic models: name length and orthographic distance. We do so in order to show that language models truly encode semantic information. In the first model, we represented each individual entity by the length of their name. In the second one, we leverage the Levenshtein distance, a popular measure of orthographic distance representing the number of letter substitutions required to transform one string into another [70]. We thus represent the orthographic properties of each name in a single value, as the average Levenshtein distance between the entity’s name and all other names in the set of stimuli.

### Encoding

#### Representational Similarity Analysis Encoding

For encoding, we used the approach proposed and validated in [71], which is based on Representation Similarity Analysis (RSA). We illustrate visually the RSA encoding procedure in Fig 2. The main advantage afforded by this methodology is that it ensures excellent performance without the need of fitting a model - which would be a concern given the small size of our dataset. RSA encoding is conceptually based on standard RSA [72], which we will here summarize shortly. Given a set of stimuli and a model that represents them in any quantitative form (numbers, vectors, etc.), the similarity between the brain responses to a given stimulus and its model representation can be measured in two steps. The first step is looking at how similar they are to all other representations in their own representational space (respectively in the brain and in the model - so-called first-order similarity). The second step consists of measuring directly the similarity between the two vectors of the pairwise similarities through second-order similarity. This approach has the advantage of providing a straightforward way to compare the representational structure of brains and models (second-order similarity), something that would be difficult given their difference in dimensionality.

**Fig 2.**
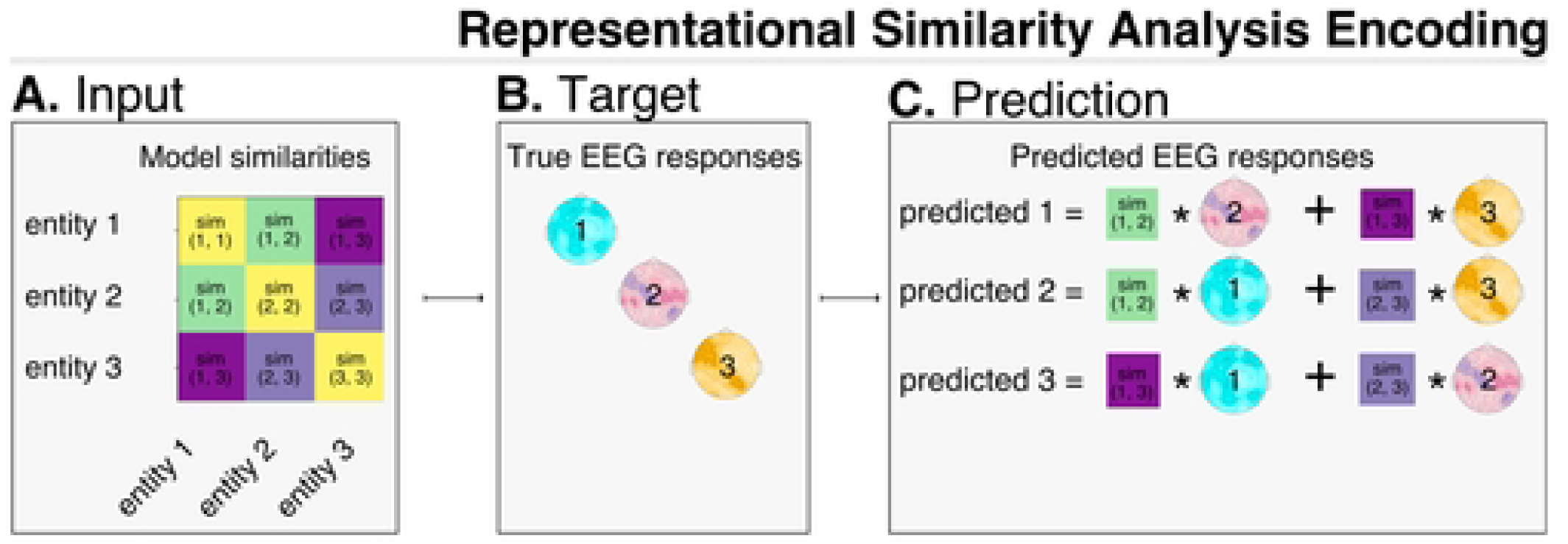
Visualization of the RSA encoding procedure. We visualize a toy example, where the whole dataset is composed of three exemplars. The input (part A) are similarities (e.g. Pearson correlations) in the model between entity representations. The targets (part B) are the corresponding EEG reponses for the same entities. Prediction happens in part C. For each entity (the test set, e.g. entity 1) its predicted EEG response corresponds to the the sum of the real evoked responses for the other entities (the train set, e.g. entity 2 and entity 3), where each response from the train set is weighted by its similarity in the model with the entity from the test set. Encoding performance is evaluated by computing the correlation between the predicted and the real response.

In RSA encoding, as in most multivariate (encoding/decoding) approaches, the data is split iteratively in a train set and a test set [50, 73]. For each train/test split, the model predicts evoked responses for the test stimuli, which are compared with the real responses. A similarity metric - in our case, Pearson correlation between predicted and real evoked responses, the most used metric used with RSA [74] - is used to evaluate how well the model captures brain activity. This procedure is repeated for all train/test split and correlations are then averaged, so as to provide an unbiased summary measurement [75].

Prediction is carried out in the following way. The evoked response to a given item from the train set is predicted as a weighted sum of the evoked responses to the stimuli in the training set. Following the original implementation, the weights to be used for the sum are the pairwise Pearson correlations between the test item and all of the training items in the model - in our case, a language model. Take for instance a toy training set composed of three individuals, for which both model representations 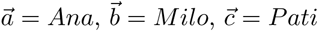, and their corresponding evoked responses *a_brain_*, *b_brain_*, *c_brain_* are available. Given a test item d = Nico, its evoked response *d_brain_* is predicted to be 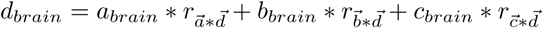, where *r* is the operation of Pearson correlation.

#### Evaluation

For evaluation we followed standard practice in encoding/decoding brain studies [50, 75]. We carried out analyses separately for each individual subject, and we then averaged results across subjects. Since for each subject we had limited data, we used a leave-two-out training/testing regime. This entails running iterated training/testing procedures, using each time as a test set one of all possible pairs of stimuli. At each iteration we recorded the Pearson correlation between each predicted test item and its true counterpart. The final correlation score was given by the average of all correlations collected by the leave-two-out train/test iterations.

#### Confound control

We controlled for name length for personally familiar people and places, since they could not be controlled *a priori* during stimulus selection, as was instead the case for famous names. To this aim, we used a cross-validated confound regression procedure that was previously validated [58]. For each train-test split, we fitted a linear regression model from the confound variable to the brain data within the train set. Then, only the residuals of the regression (also applied to the test set) entered the analyses - effectively removing from the brain data the variance that could be explained by the confound variable. We use the original python implementation provided by the authors.

#### Time-resolved encoding

In our time-resolved analyses, we ran the encoding analysis separately for every time point [50] using all of the electrodes as target for the prediction. This whole-scalp approach allows to look primarily at how distributed patterns of evoked activity develop across time. By measuring the correlation between the real brain signal and the one predicted by the model, it is possible to understand when a model captures information as it is processed in the brain. Time points where correlations are above chance with statistical significance indicate reliable encoding of such information.

#### Spatio-temporal searchlight

We were also interested in going beyond patterns of activity widely distributed across the whole scalp, and look at specific areas on the scalp where a model can explain evoked reponses. To this aim, we implemented a searchlight encoding analysis. Searchlight allows to find in a bottom-up fashion localized clusters of brain activity associated with an experimental condition, while exploiting the high sensitivity of multivariate analyses [76, 77]. In practice, searchlight consists of running the encoding analyses repeatedly across smaller clusters of electrodes on the scalp. Results are interpreted as above, just adapting the inference to the electrodes present in the cluster-correlations reliably above 0. across subjects indicate that a model can reliably predict brain activity in a given cluster at a given point in time. To reduce the computational strain, we followed previous work [78] and used spatio-temporal clusters, where multiple time points are considered for each electrode within the cluster. As in [79], we employed a temporal radius of 50ms and a spatial radius of 30mm (i.e. a cluster contains evoked activity for 100ms, for electrodes falling within a circle having a diameter of 60mm).

#### Statistical testing and corrections for multiple comparisons

We ran one-tailed statistical tests, testing the hypothesis that correlations between real and predicted scores were reliably above chance (*correlation_chance_* = 0.) across subjects, separately for each time point (time-resolved analysis) or spatio-temporal cluster (searchlight) within a time window between 0 and 800ms [52]. To control the risk of false positives, inherent when running such a high number of statistical tests, we used the Threshold-Free Cluster Enhancement (TFCE) procedure to control for multiple comparisons [50, 80, 81], implemented in MNE.

## Results

### Comparing language models

In Fig 3 we report the overall encoding results for the time-resolved and searchlight analyses, respectively using all electrodes and localized clusters of electrodes. Encoding performance is measured as Pearson correlation among the predicted and the real evoked responses; the threshold used for statistical significance is *p* < 0.05 after TFCE correction for multiple comparisons.

**Fig 3.**
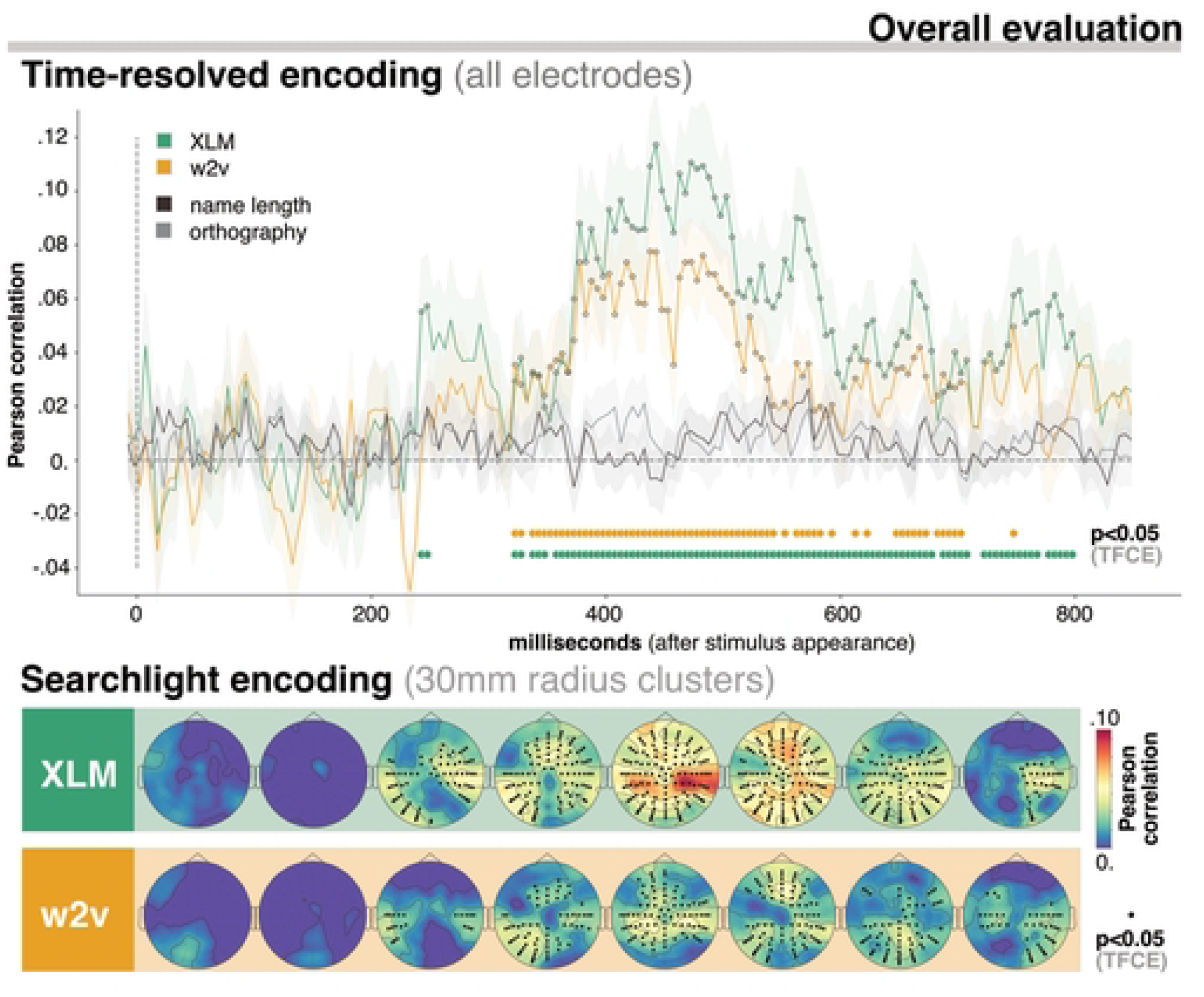
Overall encoding performance for XLM and word2vec. In the upper part of the plot we report time-resolved results obtained using all electrodes. The lower part contains searchlight results, which reflect encoding performance for smaller spatio-temporal clusters. The evaluation metric used was Pearson correlation (on the y-axis for time-resolved analysis, color-coded for searchlight). Time (time-resolved) and space-time (searchlight) points where encoding is significantly above chance (*p* < 0.05 after TFCE correction) are marked by thicker dots either at the bottom of the plot (time-resolved) or on scalp locations(searchlight). Results reported here are averaged across all entity types. They indicate overall that both XLM and word2vec capture semantic representations of individual entities in a large time window (200-800ms). In terms of spatial localization, significant performance was found in bilateral temporo-parietal and central fronto-posterior areas. We also report results for two control models (name length and orthography) for which performance was always at chance.

In the time-resolved analysis, for XLM, the contextualized language model, correlation scores were significantly above zero in a wide time window, from 250 to 800 ms (peaks between 380 and 580ms, *p* < 0.001). For the static language model, by contrast, the time window where the predicted responses correlated significantly with the real evoked potentials was shorter, between 300 and 700ms (peaks between 380 and 500ms, *p* < 0.001). Also, across all time points, correlation was consistently higher for XLM when compared to word2vec.

In searchlight, XLM provides significant correlations from 200 to 800ms. Clusters could initially (200-300ms) be found in both hemispheres, in temporo-parietal electrodes (*p* < 0.001). Encoding with XLM kept providing significant scores in bilateral electrodes up until 700ms (*p* < 0.001). After that time point, significant clusters were right lateralized, again in temporo-parietal regions (peaks in B15 to B26, *p* < 0.001). Correlations with evoked responses in central regions were significant in smaller time windows - frontally, between 300 and 600ms (*p* < 0.001); posteriorly, between 400 and 700ms (*p* < 0.001).

For searchlight encoding with word2vec, clusters where correlation was significantly above zero emerged from 200-300ms. Significant clusters for the static language model followed a similar spatial and temporal distribution as XLM. Significant correlations emerged in bilateral temporo-parietal electrodes between 200 and 800ms, in fronto-central electrodes between 300 and 600ms and in posterior-central electrodes between 400 and 700ms.

Regarding the two baseline non-semantic models (name length and orthography), correlations were never above chance. This confirms that such confounds were successfully removed from the signal through the confound removal procedure.

### XLM: encoding scores for specific types of entities

In Fig 4 we detail how the best-performing model, XLM, captures semantic information for each specific type of individual entity (personally familiar places, famous places, personally familiar people, famous people). The goal of this analysis is understanding whether our small-scale distributional information, as encoded by a contextualized language model, worked equally well for all types of entities, despite their obvious differences.

**Fig 4.**
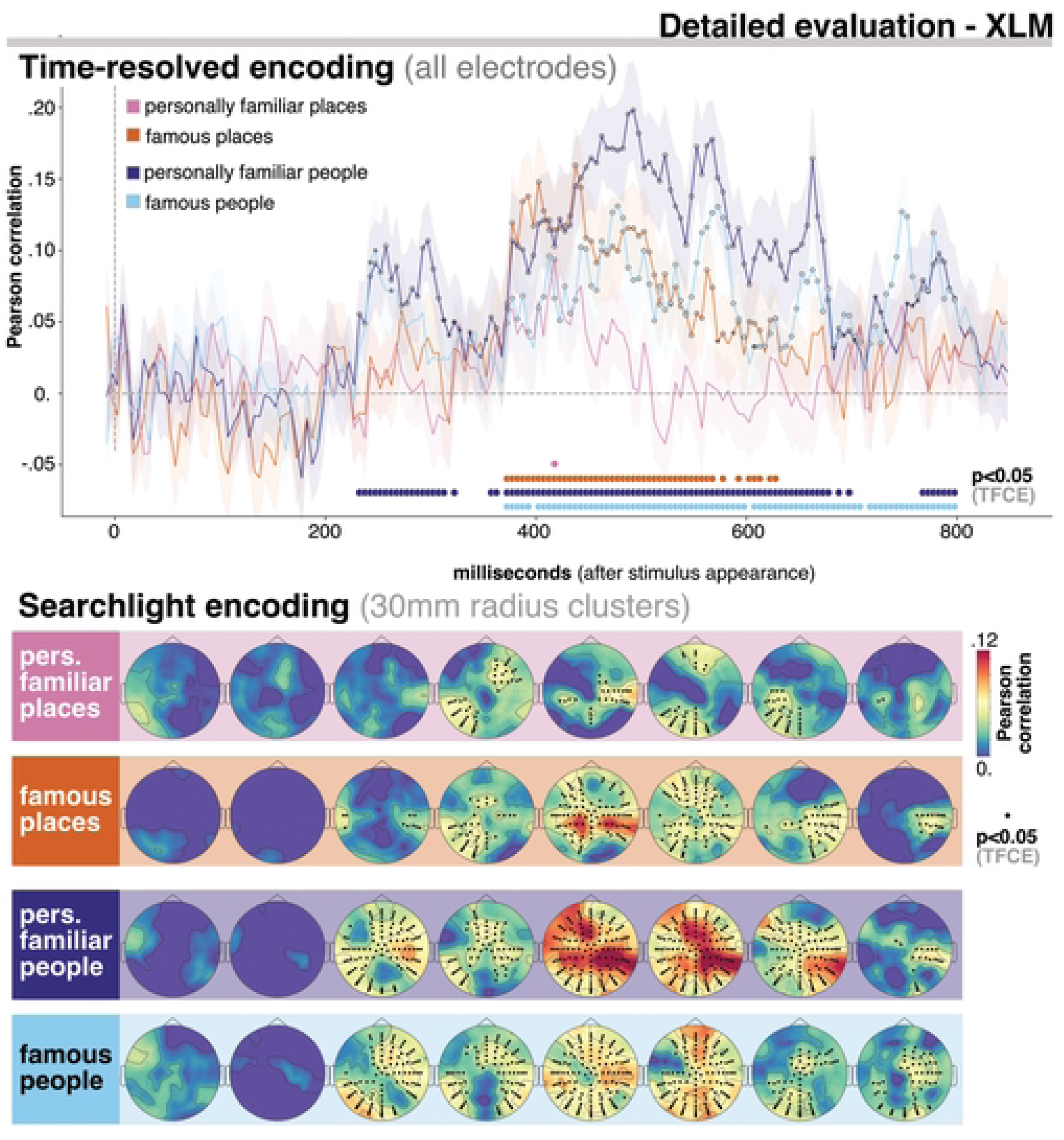
Detailed evaluation of encoding performance for XLM for all categories and levels of familiarity. In the upper part of the plot we report time-resolved results obtained using all electrodes. The lower part contains searchlight results, which reflect encoding performance for smaller spatio-temporal clusters. The evaluation metric used was Pearson correlation (on the y-axis for time-resolved analysis, color-coded for searchlight). Time (time-resolved) and space-time (searchlight) points where encoding is significantly above chance (*p* < 0.05 after TFCE correction) are marked by thicker dots either at the bottom of the plot (time-resolved) or on scalp locations(searchlight). Results reported here refer to XLM only, and are separated for each entity type, allowing direct comparisons. XLM best captured semantic processing in the brain of personally familiar people. Performance for famous people and places was, by contrast, largely similar. The toughest type of representation to capture for XLM from text were personally familiar places, as it can be seen from the lowest encoding scores overall. The spatial distribution of statistically significant scores was centered in temporo-parietal and central fronto-posterior areas.

To report scores separately for each type of entity, we repeated the encoding analyses using as test items only the stimuli belonging to the relevant category. The rationale is that performance on a test item is a quantification of the extent with which a model is able to capture the specific phenomenon of which the test item is an instance [29, 82–86].

Both time-resolved and searchlight results indicate that XLM captured in the best way personally familiar people, and in the worst way personally familiar places. For personally familiar people, correlation was significantly above zero between 200 and 800ms in a large bilateral set of electrodes, concentrated in temporo-parietal and fronto-parietal areas (*p* < 0.001). Interestingly, right-hemisphere temporo-parietal electrodes provided the highest and most consistent encoding scores overall. For personally familiar places the only significant time point in the time-resolved analysis was 0.4175ms (p = 0.04). Significant clusters of electrodes were first aligned over a bilateral temporo-parietal axis (300-500ms, *p* < 0.001), and later over a central fronto-posterior axis (500-700ms, *p* < 0.001).

Famous entities showed a less dramatic difference across categories: in the time-resolved setting, famous people could be encoded with significant correlation between 350 and 800ms (peak between 550ms and 580ms, *p* < 0.001), whereas famous places did so in the 350-650ms range (peaks between 380ms and 440ms, *p* < 0.001). Searchlight encoding revealed a clear difference in early spatial patterns (200-400ms). Semantic processing of famous people, but not of famous places, could be captured using XLM in a set of right fronto-temporal electrodes (B15 to C23, *p* < 0.001). Between 400 and 600ms, encoding performance was rather similar across the two categories - the main difference being that encoding scores for famous people were higher than for famous places. Finally, after 600ms, once again a cluster selective for famous people emerged, this time in left posterior-parietal portions of the scalp (A10 to A32, *p* < 0.001). In right temporo-parietal areas, encoding performance was similar for famous people and places.

## Discussion

### Personal memories as a window into semantic processing of individual entities

The main contribution of this work is showing that personal memories revolving around a subject’s closest people and places, collected from participants through a questionnaire, can be used to create semantic representations with language models that capture how the brain represents those very same individuals.

Our approach was motivated by the hypothesis that the way in which we talk about the world reflect our internal cognitive states [87] – and, more specifically for semantics, the idiosyncratic perspective that shapes the way in which we see the world and represent concepts [34]. Such hypothesis is supported by previous results showing that textual analyses of a speaker’s utterances can be used to uncover a speaker’s unique perspective on concepts [30, 88], their emotions [89, 90], their personality traits and mental health [91–96].

Time-wise, encoding scores were statistically significant in the 300-800ms range, with peaks at around 400ms and 700ms, which is where traditionally semantic and memory retrieval processing has been found in EEG studies – specifically, N400 [97] and late posterior complex, LPC [98].

Spatial patterns of encoding emerging from the searchlight analysis highlight the role, overall, of temporo-parietal bilateral regions of the scalp in the 400-500ms range. Such electrode clusters are situated roughly above portions of the cortex which have been shown to be crucial for the processing of individual entities by a vast literature from neuroimaging (summarized in [99]). This provides new evidence that markers of processing of proper names on the scalp seem to follow a similar trajectory as cortical activity (Fig 3, lower portion; see also [52] for similar decoding maps).

This consistent pattern of results therefore confirms our hypothesis. The distributional lexical information contained in the questionnaire answers, describing the three main components of memories for individual entities - episodic and semantic knowledge, as well as personal semantics - can be used to encode how the brain represents those individual entities.

### Using language models to encode personally familiar entities

We were able to encode responses to individual entities by exploiting, through language models, the distributional lexical information contained in the questionnaire answers. Our results thus indicate that the semantic representations obtained from language models are rich, multi-faceted models of brain processing (cfr. [11, 100]) and that, following an appropriate methodology, they can be combined to create vectors for entities that do not appear in their pre-training data [27, 30]. The questionnaire was composed of nine questions only, to which participants had to answer to with open-ended natural sentences - which by Natural Language Processing (NLP) standards is extremely small [24].

This result is all the more surprising given the small size of the textual data, and the fact that participants produced answers freely, without having to stick to categories or dimensions defined a priori by the experimenter as was the case in previous neuroimaging studies [31–33].

A key role in making the most of such reduced textual data was played by a dedicated vector creation methodology (see Section). We created original semantic representations for personally familiar individual entities from the unconstrained questionnaire answers, building upon the previously-acquired distributional lexical knowledge contained in language models. In other words, through language models we could model semantic knowledge of people and places as idiosyncratic combinations of pieces of generic knowledge (the pre-trained word vectors), shaped and guided by autobiographic experiences (the person-specific words contained in the answers to the questionnaires).

In this respect, XLM, the contextualized language model, provides consistently superior performance, when compared to word2vec, a static language model, at capturing semantic processing in the brain (Fig 3). Notice that we matched the vector extraction methodology, thus levelling as much as possible the differences between the two models. Two remaining factors differentiate static and contextual models. The first one the size (see Section). The second one is that contextualized models are able to adapt their vector representations to specific linguistic contexts, capturing fine-grained contextual shifts in meaning, whereas static models cannot [61]. These two characteristics, therefore, seem to be crucial in order to capture the semantic information revolving around an individual entity, which is an extremely idiosyncratic and unique mix of features and memories [1, 2]. Also, this evidence dovetails with previous results in the literature indicating that such superior performance holds also for generic concepts and famous individual entities [29, 101–103].

Our findings would seem to be compatible with so-called **descriptivism**, a view on proper names and individual entities that proposes that the meaning of a proper name can be in fact equated to a set of sentences describing that entity (in our case, declarative memories) [104, 105]. However, one could also argue, as pointed out by [99], that descriptions (and by extension, declarative knowledge) fail at capturing all semantic information associated with cognitive representations of individual entities, namely not considering socio-emotional dimensions. We believe that it is precisely our choice of modelling memories (i.e. descriptions) through language models that allows to address this concern. Language models have been consistently shown to capture both social and emotional dimensions of word meaning from lexical distributional information [11, 13, 14]. Therefore, our view is that although social and emotional semantic dimensions of individual entities can be studied in isolation (cfr. [31–33]), they are latent in declarative memories (descriptions) and can be therefore be captured in the mixture of semantic information and dimensions present in language models [100]. In this sense, descriptions are a richer source of semantic information about individual entities than it may appear at first sight: they not only convey their propositional content, but also the broader semantic information hidden in the words that make them up.

### Individual entities

Detailed patterns of encoding performance can be compared in terms of scores and spatial location (Fig 4) across semantic categories (people and places) and levels of familiarity (famous and personally familiar). This allows to obtain insights with respect to the way in which the brain represents individual entities.

First of all, it is important to notice that, as experimental stimuli, we used names, instead of images - which is by contrast the most common choice when looking at concepts in the brain [51], possibly because they afford higher encoding/decoding performance [18, 52, 106, 107]. The reason why we chose to use names in our experiment is that this would allow us to elicit semantic processing of people and places in the brain not biased towards any sensory modality. For instance, had we used images as stimuli, we would have had two types of non-semantic, visual confounds. First, images clearly differ in terms of visual properties across people and places and therefore evoke strongly distinct responses [108, 109]. Secondly, using a picture elicits brain activity associated with that specific instance of the picture and its low-level visual features [110, 111].

Overall, we found that the semantics of people, both personally familiar and famous, could be captured using distributional lexical information representing declarative memories (in its three components - semantic, episodic and personal) - despite the uniqueness of the semantic information for each individual [1–3, 99]. For places, encoding patterns differed across levels of familiarity. Famous places showed similar encoding scores as famous people. Personally familiar places, by contrast, had the lowest encoding scores overall, suggesting that they are particularly challenging for models based on distributional semantics.

Searchlight encoding, which provides a window on focal activity on the scalp, highlights both commonalities and differences for the various types of individual entities.

When looking at the commonalities, a core set of spatio-temporal clusters shared across categories and levels of familiarity emerges, indicating general correlates of semantic processing of individual entities. Encoding was significant for all type of individual entities in the range between 300 and 700ms, first in temporo-parietal bilateral electrodes (300-500ms), then fronto- and posterior-central electrodes (500-700ms). The former are typically associated with general semantic processing in the N400 time range [51, 97], whereas the latter could be explained as an involvement of both Default Mode Network areas, which are activated by episodic memory and social processing [112–115], and of posterior visual areas relevant for processing mental imagery [116, 117].

Turning to the differences, the earliest time range where correlations were significant (200-300ms) revealed an interesting pattern: a stronger presence of information related to people as opposed to places in the fronto-temporal right hemisphere (Fig 4, lower portion). This can be connected to results coming from the neuroimaging literature, where it has been shown that the right hemisphere is crucial specifically for person identity processing [118–122].

## Conclusion

In this work we have shown that it is possible to capture semantic knowledge about personally familiar individual entities by extracting semantic representations from subject-specific memories using language models. Importantly, we also demonstrated that similar performance could be obtained also for a matched set of famous entities, thus proving the solidity of our approach. The results of our multivariate encoding analyses indicate that entity-specific information emerges in a time window usually associated with semantic processing, between 200-800ms. Also, this seems to be the case especially in bilateral temporo-parietal regions, which converges with neuroimaging studies. Overall, our results exploit cutting-edge models in AI to provide a window into extremely fine-grained, subject-specific semantic knowledge as it is processed in the brain. We hope that this will motivate future work aiming at the investigation of individual uniqueness in semantic processing.

## Acknowledgments

We would like to thank prof. Davide Crepaldi, head of the Language, Learning and Reading lab at SISSA, who provided the facilities for collecting the EEG data while the first author was visiting, and Marjina Bellida for helping out with subject preparation procedures.

## Footnotes

1 https://azure.microsoft.com/en-us/products/cognitive-services/speech-to-text

## References

1. Cohen G, Burke DM. Memory for proper names: A review. Memory. 1993;1(4):249–263.

2. Kaminski J, Bowren Jr M, Manzel K, Tranel D. Neural correlates of recognition and naming of famous persons and landmarks: A special role for the left anterior temporal lobe. In: Handbook of Clinical Neurology. vol. 187. Elsevier; 2022. p. 303–317.

3. Semenza C. Proper names and personal identity. Handbook of Clinical Neurology. 2022;187:287–302.

4. Tulving E. Episodic and semantic memory. Organization of Memory. 1972; p. 382–403.

5. Tulving E. Episodic memory: From mind to brain. Annual review of psychology. 2002;53(1):1–25.

6. Yee E, Chrysikou EG, Thompson-Schill SL. Semantic Memory 17. The Oxford Handbook of Cognitive Neuroscience: Volume 1: Core Topics. 2013; p. 353.

7. Renoult L, Davidson PS, Palombo DJ, Moscovitch M, Levine B. Personal semantics: at the crossroads of semantic and episodic memory. Trends in cognitive sciences. 2012;16(11):550–558.

8. Eichenbaum H. Declarative memory: Insights from cognitive neurobiology. Annual review of psychology. 1997;48(1):547–572.

9. Wittgenstein L. Philosophical investigations. Philosophische Untersuchungen. Macmillan; 1953.

10. Harris ZS. Distributional structure. Word. 1954;10(2-3):146–162.

11. Hollis G, Westbury C. The principals of meaning: Extracting semantic dimensions from co-occurrence models of semantics. Psychonomic bulletin & review. 2016;23:1744–1756.

12. Hollis G, Westbury C, Lefsrud L. Extrapolating human judgments from skip-gram vector representations of word meaning. Quarterly Journal of Experimental Psychology. 2017;70(8):1603–1619.

13. Utsumi A. Exploring what is encoded in distributional word vectors: A neurobiologically motivated analysis. Cognitive Science. 2020;44(6):e12844.

14. Chersoni E, Santus E, Huang CR, Lenci A. Decoding word embeddings with brain-based semantic features. Computational Linguistics. 2021;47(3):663–698.

15. Mandera P, Keuleers E, Brysbaert M. Explaining human performance in psycholinguistic tasks with models of semantic similarity based on prediction and counting: A review and empirical validation. Journal of Memory and Language. 2017;92:57–78.

16. Wingfield C, Connell L. Understanding the role of linguistic distributional knowledge in cognition. Language, Cognition and Neuroscience. 2022;37(10):1220–1270.

17. Mitchell TM, Shinkareva SV, Carlson A, Chang KM, Malave VL, Mason RA, et al. Predicting human brain activity associated with the meanings of nouns. science. 2008;320(5880):1191–1195.

18. Pereira F, Lou B, Pritchett B, Ritter S, Gershman SJ, Kanwisher N, et al. Toward a universal decoder of linguistic meaning from brain activation. Nature communications. 2018;9(1):963.

19. Goldstein A, Zada Z, Buchnik E, Schain M, Price A, Aubrey B, et al. Shared computational principles for language processing in humans and deep language models. Nature neuroscience. 2022;25(3):369–380.

20. Hale JT, Campanelli L, Li J, Bhattasali S, Pallier C, Brennan JR. Neurocomputational models of language processing. Annual Review of Linguistics. 2022;8:427–446.

21. Recchia G, Jones MN. More data trumps smarter algorithms: Comparing pointwise mutual information with latent semantic analysis. Behavior research methods. 2009;41:647–656.

22. Sahlgren M, Lenci A. The Effects of Data Size and Frequency Range on Distributional Semantic Models. In: Proceedings of the 2016 Conference on Empirical Methods in Natural Language Processing; 2016. p. 975–980.

23. Günther F, Rinaldi L, Marelli M. Vector-space models of semantic representation from a cognitive perspective: A discussion of common misconceptions. Perspectives on Psychological Science. 2019;14(6):1006–1033.

24. Ruzzetti ES, Ranaldi L, Mastromattei M, Fallucchi F, Scarpato N, Zanzotto FM. Lacking the Embedding of a Word? Look it up into a Traditional Dictionary. In: Findings of the Association for Computational Linguistics: ACL 2022; 2022. p. 2651–2662.

25. Yu W, Zhu C, Fang Y, Yu D, Wang S, Xu Y, et al. Dict-BERT: Enhancing Language Model Pre-training with Dictionary. In: Findings of the Association for Computational Linguistics: ACL 2022; 2022. p. 1907–1918.

26. Asr FT, Willits JA, Jones MN. Comparing Predictive and Co-occurrence Based Models of Lexical Semantics Trained on Child-directed Speech. In: 38th Annual Meeting of the Cognitive Science Society: Recognizing and Representing Events, CogSci 2016. The Cognitive Science Society; 2016. p. 1092–1097.

27. Herbelot A, Baroni M. High-risk learning: acquiring new word vectors from tiny data. In: Proceedings of the 2017 Conference on Empirical Methods in Natural Language Processing; 2017. p. 304–309.

28. Schick T, Schütze H. Rare words: A major problem for contextualized embeddings and how to fix it by attentive mimicking. In: Proceedings of the AAAI Conference on Artificial Intelligence. vol. 34; 2020. p. 8766–8774.

29. Bruera A, Poesio M. Exploring the representations of individual entities in the brain combining EEG and distributional semantics. Frontiers in Artificial Intelligence. 2022; p. 25.

30. Anderson AJ, McDermott K, Rooks B, Heffner KL, Dodell-Feder D, Lin FV. Decoding individual identity from brain activity elicited in imagining common experiences. Nature communications. 2020;11(1):5916.

31. Thornton MA, Weaverdyck ME, Tamir DI. The brain represents people as the mental states they habitually experience. Nature communications. 2019;10(1):2291.

32. Peer M, Hayman M, Tamir B, Arzy S. Brain coding of social network structure. Journal of Neuroscience. 2021;41(22):4897–4909.

33. Ron Y, Dafni-Merom A, Saadon-Grosman N, Roseman M, Elias U, Arzy S. Brain system for social categorization by narrative roles. Journal of Neuroscience. 2022;42(26):5246–5253.

34. Charest I, Kriegeskorte N. The brain of the beholder: honouring individual representational idiosyncrasies. Language, Cognition and Neuroscience. 2015;30(4):367–379.

35. De Haas B, Iakovidis AL, Schwarzkopf DS, Gegenfurtner KR. Individual differences in visual salience vary along semantic dimensions. Proceedings of the National Academy of Sciences. 2019;116(24):11687–11692.

36. Levine SM, Schwarzbach JV. Individualizing representational similarity analysis. Frontiers in psychiatry. 2021;12:729457.

37. Chen M, Chu Z, Chen Y, Stratos K, Gimpel K. EntEval: A Holistic Evaluation Benchmark for Entity Representations. In: Proceedings of the 2019 Conference on Empirical Methods in Natural Language Processing and the 9th International Joint Conference on Natural Language Processing (EMNLP-IJCNLP); 2019. p. 421–433.

38. Westera M, Gupta A, Boleda G, Padó S. Distributional models of category concepts based on names of category members. Cognitive Science. 2021;45(9):e13029.

39. Mikolov T, Sutskever I, Chen K, Corrado GS, Dean J. Distributed representations of words and phrases and their compositionality. Advances in neural information processing systems. 2013;26.

40. Conneau A, Khandelwal K, Goyal N, Chaudhary V, Wenzek G, Guzmán F, et al. Unsupervised Cross-lingual Representation Learning at Scale. In: Proceedings of the 58th Annual Meeting of the Association for Computational Linguistics; 2020. p. 8440–8451.

41. Vaswani A, Shazeer N, Parmar N, Uszkoreit J, Jones L, Gomez AN, et al. Attention is all you need. Advances in neural information processing systems. 2017;30.

42. Moore V, Valentine T. The Effects Of Age Of Acquisition In Processing Famous Faces And Names: Exploring The Locus And Proposing A Mechanism. In: Proceedings of the Twenty First Annual Conference of the Cognitive Science Society. Psychology Press; 2020. p. 416–421.

43. Paivio A, Yuille JC, Madigan SA. Concreteness, imagery, and meaningfulness values for 925 nouns. Journal of experimental psychology. 1968;76(1p2):1.

44. Rofes A, Zakariás L, Ceder K, Lind M, Johansson MB, De Aguiar V, et al. Imageability ratings across languages. Behavior Research Methods. 2018;50:1187–1197. doi:10.3758/s13428-017-0936-0.

45. Morton NW, Zippi EL, Noh SM, Preston AR. Semantic knowledge of famous people and places is represented in hippocampus and distinct cortical networks. Journal of Neuroscience. 2021;41(12):2762–2779.

46. Zhou WX, Sornette D, Hill RA, Dunbar RI. Discrete hierarchical organization of social group sizes. Proceedings of the Royal Society B: Biological Sciences. 2005;272(1561):439–444.

47. Hill RA, Dunbar RI. Social network size in humans. Human nature. 2003;14(1):53–72.

48. Bruera A, Poesio M. EEG searchlight decoding reveals person- and place-specific responses for semantic category and familiarity; 2023.

49. Boudewyn MA, Luck SJ, Farrens JL, Kappenman ES. How many trials does it take to get a significant ERP effect? It depends. Psychophysiology. 2018;55(6):e13049. doi:10.1111/psyp.13049.

50. Grootswagers T, Wardle SG, Carlson TA. Decoding dynamic brain patterns from evoked responses: A tutorial on multivariate pattern analysis applied to time series neuroimaging data. Journal of cognitive neuroscience. 2017;29(4):677–697.

51. Rybář M, Poli R, Daly I. Decoding of semantic categories of imagined concepts of animals and tools in fNIRS. Journal of Neural Engineering. 2021;18(4):046035.

52. Leonardelli E, Fait E, Fairhall SL. Temporal dynamics of access to amodal representations of category-level conceptual information. Scientific reports. 2019;9(1):239.

53. Gramfort A, Luessi M, Larson E, Engemann DA, Strohmeier D, Brodbeck C, et al. MEG and EEG data analysis with MNE-Python. Frontiers in neuroscience. 2013; p. 267.

54. Jas M, Larson E, Engemann DA, Leppäkangas J, Taulu S, Hämäläinen M, et al. A reproducible MEG/EEG group study with the MNE software: recommendations, quality assessments, and good practices. Frontiers in neuroscience. 2018;12:530.

55. Luck SJ. An introduction to the event-related potential technique. MIT press; 2014.

56. Jas M, Engemann DA, Bekhti Y, Raimondo F, Gramfort A. Autoreject: Automated artifact rejection for MEG and EEG data. NeuroImage. 2017;159:417–429.

57. Todd MT, Nystrom LE, Cohen JD. Confounds in multivariate pattern analysis: theory and rule representation case study. Neuroimage. 2013;77:157–165.

58. Snoek L, Miletić S, Scholte HS. How to control for confounds in decoding analyses of neuroimaging data. Neuroimage. 2019;184:741–760.

59. Bernardi R, Boleda G, Fernández R, Paperno D. Distributional semantics in use. In: Proceedings of the First Workshop on Linking Computational Models of Lexical, Sentential and Discourse-level Semantics; 2015. p. 95–101.

60. Kim N, Patel R, Poliak A, Xia P, Wang A, McCoy T, et al. Probing What Different NLP Tasks Teach Machines about Function Word Comprehension. In: Proceedings of the Eighth Joint Conference on Lexical and Computational Semantics (* SEM 2019); 2019. p. 235–249.

61. Apidianaki M. From Word Types to Tokens and Back: A Survey of Approaches to Word Meaning Representation and Interpretation. Computational Linguistics. 2022; p. 1–60.

62. Yee E, Jones MN, McRae K. Semantic memory. The Stevens’ handbook of experimental psychology and cognitive neuroscience. 2018;3.

63. Lenci A, Sahlgren M, Jeuniaux P, Cuba Gyllensten A, Miliani M. A comparative evaluation and analysis of three generations of Distributional Semantic Models. Language resources and evaluation. 2022;56(4):1269–1313.

64. Bender EM, Gebru T, McMillan-Major A, Shmitchell S. On the Dangers of Stochastic Parrots: Can Language Models Be Too Big? In: Proceedings of the 2021 ACM conference on fairness, accountability, and transparency; 2021. p. 610–623.

65. Vulić I, Ponti EM, Litschko R, Glavaš G, Korhonen A. Probing Pretrained Language Models for Lexical Semantics. In: Proceedings of the 2020 Conference on Empirical Methods in Natural Language Processing (EMNLP); 2020. p. 7222–7240.

66. Pires T, Schlinger E, Garrette D. How Multilingual is Multilingual BERT? In: Proceedings of the 57th Annual Meeting of the Association for Computational Linguistics; 2019. p. 4996–5001.

67. Wu S, Dredze M. Are All Languages Created Equal in Multilingual BERT? In: Proceedings of the 5th Workshop on Representation Learning for NLP; 2020. p. 120–130.

68. Mass Y, Roitman H, Erera S, Rivlin O, Weiner B, Konopnicki D. A study of bert for non-factoid question-answering under passage length constraints. arXiv preprint arXiv:190806780. 2019;.

69. Rudnicka K. Variation of sentence length across time and genre. Diachronic corpora, genre, and language change. 2018; p. 220–240.

70. Yarkoni T, Balota D, Yap M. Moving beyond Coltheart’s N: A new measure of orthographic similarity. Psychonomic bulletin & review. 2008;15(5):971–979.

71. Anderson AJ, Zinszer BD, Raizada RD. Representational similarity encoding for fMRI: Pattern-based synthesis to predict brain activity using stimulus-model-similarities. NeuroImage. 2016;128:44–53.

72. Kriegeskorte N, Mur M, Bandettini PA. Representational similarity analysis-connecting the branches of systems neuroscience. Frontiers in systems neuroscience. 2008; p. 4.

73. Walther A, Nili H, Ejaz N, Alink A, Kriegeskorte N, Diedrichsen J. Reliability of dissimilarity measures for multi-voxel pattern analysis. Neuroimage. 2016;137:188–200.

74. Nili H, Wingfield C, Walther A, Su L, Marslen-Wilson W, Kriegeskorte N. A toolbox for representational similarity analysis. PLoS computational biology. 2014;10(4):e1003553.

75. Varoquaux G, Raamana PR, Engemann DA, Hoyos-Idrobo A, Schwartz Y, Thirion B. Assessing and tuning brain decoders: cross-validation, caveats, and guidelines. NeuroImage. 2017;145:166–179.

76. Etzel JA, Zacks JM, Braver TS. Searchlight analysis: promise, pitfalls, and potential. Neuroimage. 2013;78:261–269.

77. Kriegeskorte N, Goebel R, Bandettini P. Information-based functional brain mapping. Proceedings of the National Academy of Sciences. 2006;103(10):3863–3868.

78. Su L, Fonteneau E, Marslen-Wilson W, Kriegeskorte N. Spatiotemporal searchlight representational similarity analysis in EMEG source space In: 2012 Second International Workshop on Pattern Recognition in NeuroImaging; 2012.

79. Collins E, Robinson AK, Behrmann M. Distinct neural processes for the perception of familiar versus unfamiliar faces along the visual hierarchy revealed by EEG. NeuroImage. 2018;181:120–131.

80. Smith SM, Nichols TE. Threshold-free cluster enhancement: addressing problems of smoothing, threshold dependence and localisation in cluster inference. Neuroimage. 2009;44(1):83–98.

81. Latinus M, Nichols T, Rousselet G. Cluster-based computational methods for mass univariate analyses of event-related brain potentials/fields: A simulation study. Journal of neuroscience methods. 2015;250:85–93.

82. Kaplan JT, Man K, Greening SG. Multivariate cross-classification: applying machine learning techniques to characterize abstraction in neural representations. Frontiers in human neuroscience. 2015;9:151.

83. Lake B, Baroni M. Generalization without systematicity: On the compositional skills of sequence-to-sequence recurrent networks. In: International conference on machine learning. PMLR; 2018. p. 2873–2882.

84. Gorman K, Bedrick S. We need to talk about standard splits. In: Proceedings of the 57th annual meeting of the association for computational linguistics; 2019. p. 2786–2791.

85. Elangovan A, He J, Verspoor K. Memorization vs. Generalization: Quantifying Data Leakage in NLP Performance Evaluation. In: Proceedings of the 16th Conference of the European Chapter of the Association for Computational Linguistics: Main Volume; 2021. p. 1325–1335.

86. Chyzhyk D, Varoquaux G, Milham M, Thirion B. How to remove or control confounds in predictive models, with applications to brain biomarkers. GigaScience. 2022;11.

87. Eichstaedt JC, Kern ML, Yaden DB, Giorgi S, Park G, Hagan CA, et al. Closed-and open-vocabulary approaches to text analysis: A review, quantitative comparison, and recommendations. Psychological Methods. 2021;26(4):398.

88. Herbelot A, QasemiZadeh B. You and me… in a vector space: modelling individual speakers with distributional semantics. In: Proceedings of the Fifth Joint Conference on Lexical and Computational Semantics; 2016. p. 179–188.

89. Strapparava C, Mihalcea R. Learning to identify emotions in text. In: Proceedings of the 2008 ACM symposium on Applied computing; 2008. p. 1556–1560.

90. Sailunaz K, Dhaliwal M, Rokne J, Alhajj R. Emotion detection from text and speech: a survey. Social Network Analysis and Mining. 2018;8:1–26.

91. Pennebaker JW, Graybeal A. Patterns of natural language use: Disclosure, personality, and social integration. Current Directions in Psychological Science. 2001;10(3):90–93.

92. Calvo RA, Milne DN, Hussain MS, Christensen H. Natural language processing in mental health applications using non-clinical texts. Natural Language Engineering. 2017;23(5):649–685.

93. Corcoran CM, Carrillo F, Fernández-Slezak D, Bedi G, Klim C, Javitt DC, et al. Prediction of psychosis across protocols and risk cohorts using automated language analysis. World Psychiatry. 2018;17(1):67–75.

94. Mota NB, Weissheimer J, Ribeiro M, De Paiva M, Avilla-Souza J, Simabucuru G, et al. Dreaming during the Covid-19 pandemic: Computational assessment of dream reports reveals mental suffering related to fear of contagion. PloS one. 2020;15(11):e0242903.

95. Sarzynska-Wawer J, Wawer A, Pawlak A, Szymanowska J, Stefaniak I, Jarkiewicz M, et al. Detecting formal thought disorder by deep contextualized word representations. Psychiatry Research. 2021;304:114135.

96. Grimmer J, Roberts ME, Stewart BM. Text as data: A new framework for machine learning and the social sciences. Princeton University Press; 2022.

97. Kutas M, Federmeier KD. Thirty years and counting: finding meaning in the N400 component of the event-related brain potential (ERP). Annual review of psychology. 2011;62:621–647.

98. Wilding EL, Ranganath C. Electrophysiological Correlates of Episodic Memory Processes. The Oxford Handbook of Event-Related Potential Components. 2011; p. 373.

99. O’Rourke T, de Diego Balaguer R. Names and their meanings: A dual-process account of proper-name encoding and retrieval. Neuroscience & Biobehavioral Reviews. 2020;108:308–321.

100. Grand G, Blank IA, Pereira F, Fedorenko E. Semantic projection recovers rich human knowledge of multiple object features from word embeddings. Nature human behaviour. 2022;6(7):975–987.

101. Jain S, Huth A. Incorporating context into language encoding models for fMRI. Advances in neural information processing systems. 2018;31.

102. Jat S, Tang H, Talukdar P, Mitchell T. Relating Simple Sentence Representations in Deep Neural Networks and the Brain. In: Proceedings of the 57th Annual Meeting of the Association for Computational Linguistics; 2019. p. 5137–5154.

103. Schrimpf M, Blank IA, Tuckute G, Kauf C, Hosseini EA, Kanwisher N, et al. The neural architecture of language: Integrative modeling converges on predictive processing. Proceedings of the National Academy of Sciences. 2021;118(45):e2105646118.

104. Russell B. Knowledge by acquaintance and knowledge by description. In: Proceedings of the Aristotelian society. vol. 11. JSTOR; 1910. p. 108–128.

105. Searle JR. Proper names. Mind. 1958;67(266):166–173.

106. Simanova I, Van Gerven M, Oostenveld R, Hagoort P. Identifying object categories from event-related EEG: toward decoding of conceptual representations. PloS one. 2010;5(12):e14465.

107. Shinkareva SV, Malave VL, Mason RA, Mitchell TM, Just MA. Commonality of neural representations of words and pictures. Neuroimage. 2011;54(3):2418–2425.

108. Rousselet GA, Husk JS, Bennett PJ, Sekuler AB. Time course and robustness of ERP object and face differences. Journal of vision. 2008;8(12):3–3.

109. Rossion B. Understanding face perception by means of human electrophysiology. Trends in cognitive sciences. 2014;18(6):310–318.

110. Just MA, Cherkassky VL, Aryal S, Mitchell TM. A neurosemantic theory of concrete noun representation based on the underlying brain codes. PloS one. 2010;5(1):e8622.

111. Simanova I, Hagoort P, Oostenveld R, Van Gerven MA. Modality-independent decoding of semantic information from the human brain. Cerebral cortex. 2014;24(2):426–434.

112. Raichle ME. The brain’s default mode network. Annual review of neuroscience. 2015;38:433–447.

113. Campbell A, Louw R, Michniak E, Tanaka JW. Identity-specific neural responses to three categories of face familiarity (own, friend, stranger) using fast periodic visual stimulation. Neuropsychologia. 2020;141:107415.

114. Smallwood J, Bernhardt BC, Leech R, Bzdok D, Jefferies E, Margulies DS. The default mode network in cognition: a topographical perspective. Nature reviews neuroscience. 2021;22(8):503–513.

115. Kaefer K, Stella F, McNaughton BL, Battaglia FP. Replay, the default mode network and the cascaded memory systems model. Nature Reviews Neuroscience. 2022;23(10):628–640.

116. Reddy L, Tsuchiya N, Serre T. Reading the mind’s eye: decoding category information during mental imagery. Neuroimage. 2010;50(2):818–825.

117. Xie S, Kaiser D, Cichy RM. Visual imagery and perception share neural representations in the alpha frequency band. Current Biology. 2020;30(13):2621–2627.

118. Gainotti G. Different patterns of famous people recognition disorders in patients with right and left anterior temporal lesions: a systematic review. Neuropsychologia. 2007;45(8):1591–1607.

119. Gainotti G. Implications of recent findings for current cognitive models of familiar people recognition. Neuropsychologia. 2015;77:279–287.

120. Borghesani V, Narvid J, Battistella G, Shwe W, Watson C, Binney RJ, et al. “Looks familiar, but I do not know who she is”: The role of the anterior right temporal lobe in famous face recognition. Cortex. 2019;115:72–85.

121. Pisoni A, Sperandeo PR, Lauro LJR, Papagno C. The role of the left and right anterior temporal poles in people naming and recognition. Neuroscience. 2020;440:175–185.

122. Desai RH, Tadimeti U, Riccardi N. Proper and common names in the semantic system. Brain Structure and Function. 2022; p. 1–16.

